# Early transmission of sensitive strain slows down emergence of drug resistance in *Plasmodium vivax*

**DOI:** 10.1101/603597

**Authors:** Mario J.C. Ayala, Daniel A.M. Villela

## Abstract

The spread of drug resistance of *Plasmodium falciparum* and *Plasmodium vivax* parasites is a challenge towards malaria elimination.*P. falciparum* has shown an early and severe drug resistance in comparison to *P. vivax* in various countries. In fact, these *Plasmodium* species differ in their life cycle and treatment in various factors: development and duration of sexual parasite forms differ, symptoms severity are unequal, relapses present only in *P. vivax* cases, and the Artemisinin-based combination therapy (ACT) is only mandatory in all *P. falciparum* cases. We compared the spread of drug resistance for both species through two compartmental models using ordinary differential equations. The model structure describes how sensitive and resistant parasite strains infect a human population treated with antimalarials. We found that the early transmission before treatment and the low effectiveness of drug coverage support the prevalence of sensitive parasites delaying the emergence of resistant *P. vivax*. These results imply that earlier attention of symptomatic and reservoirs of *P. vivax* accelerates the spread of drug resistance as *P. falciparum*.

## Introduction

Spread of drug resistance of *Plasmodium falciparum* and *Plasmodium vivax* parasites challenges malaria programmes towards elimination, as previous studies have confirmed the spread of drug resistance to the first-line drugs: chloroquine (CQ) and artemisinin-based combination therapies (ACTs) [1–6]. Nevertheless, resistant parasites have emerged in different geographical and temporal patterns around the endemic regions on the world [5]. Thus, efficient malaria programs require considering the variations in treatment regimens and environmental conditions.

Apart from external conditions, *P. falciparum* and *P. vivax* parasites differ in their life cycle and treatment exhibiting distinct patterns in drug resistance. *P. falciparum* has shown an early and severe drug resistance in comparison to *P. vivax*. Currently, chloroquine (CQ) remains as the first-line treatment against *P. vivax* and ACTs to *P. falciparum* due to the reports of CQ resistance in *P. falciparum* dating from 1950 [2, 3, 7]. On the other hand, these *Plasmodium* species differ in various factors: relapses only in *P. vivax* cases, development and duration of sexual parasite forms, symptoms severity and host immunity response [6, 8, 9]. Therefore, these species diverge and require specific studies in order to understand and compare determinants in their particular evolution of drug resistance.

Although most previous studies focused on *P. falciparum* resistance, a few works have compared the evolution of drug resistance between both species [5, 8, 10]. Most of previous studies evaluated *P. falciparum*-resistance factors such as cost-resistance, selection after treatment, transmission of resistant parasites, epidemiological factors, asymptomatic infections and treatment regimens [11–22]. Even though *P. vivax* caused the 74% of malaria cases in the Americas and the 37% of cases in the Asian Southwest, the impact of drug resistant in parasite prevalence still remains underestimate [23].

Our aim is to compare the emergence and spread of *P. vivax* and *P. falciparum* drug resistance taking into account their particular life cycles. Here, we developed compartmental models for both *P. vivax* and *P. falciparum* illustrating the emergence and transmission of resistant parasites under different treatment regimens and epidemiological conditions in human population level. Our approach reveals the impact in drug resistance of both species filling in the gap of knowledge about *P. vivax* resistance.

## Materials and methods

We developed mathematical models for both *Plasmodium vivax* and *Plasmodium falciparum* using ordinary differential equations (ODE) to represent the transmission of two strains: sensitive and resistant. These models are based on the well-known Ross-Macdonald structure that measure human and mosquito populations dividing by susceptible and infected individuals [24]. Additionally, we implemented a post-treatment state in humans and we also divided infected states by sensitive and resistant. The next subsections expand model features and differences between *P. vivax* and *P. falciparum* modeling.

### *P. falciparum* model

This model outlines *P. falciparum* transmission in five human and three mosquito states: susceptible humans *S*_*h*_, infected humans by sensitive strain *I*_*fs*_, post-treatment humans after sensitive infection *P*_*fs*_, infected humans by resistant strain *I*_*fr*_, post-treatment humans after resistant infection *P*_*fr*_, susceptible mosquitoes *S*_*m*_, infected mosquitoes by sensitive strain *I*_*mfs*_, and infected mosquitoes by resistant strain *I*_*mfr*_. Infected and post-treatment humans can infect susceptible mosquitoes and then, they become susceptible again (see Fig 1). On the other hand, infected mosquitoes remain in this state until their death due to their the short life expectancy. The equations from Eq 1 to Eq 8 represent the measure per state; table 1 illustrates model parameters.

**Table 1.**
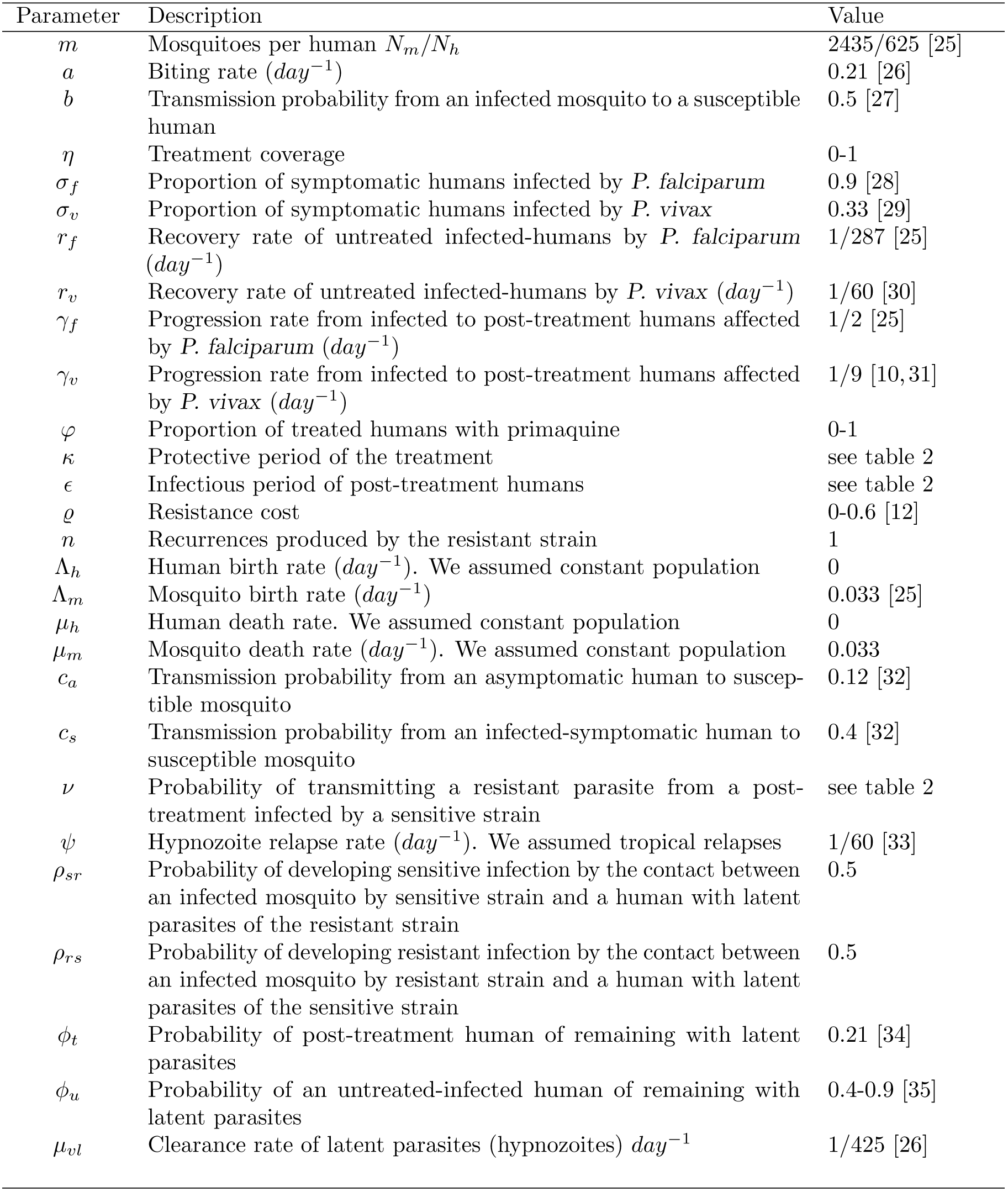
Model parameters.

**Fig 1.**
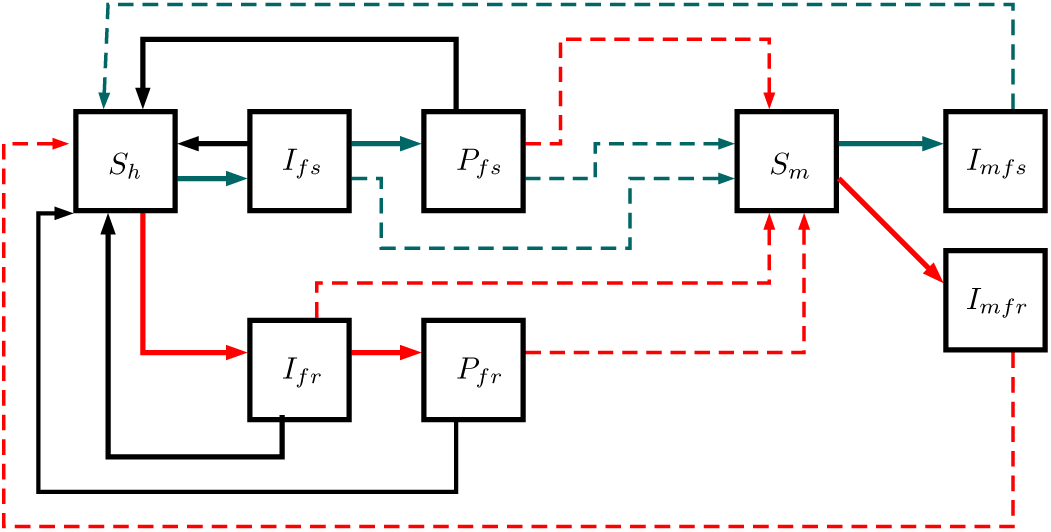
*P. Falciparum* model. This structure illustrates the transmission in five human and three mosquito states: susceptible humans *S*_*h*_, infected humans by sensitive strain *I*_*fs*_, post-treatment humans after sensitive infection *P*_*fs*_, infected humans by resistant strain *I*_*fr*_, post-treatment humans after resistant infection *P*_*fr*_, susceptible mosquitoes *S*_*m*_, infected mosquitoes by sensitive strain *I*_*mfs*_, and infected mosquitoes by resistant strain *I*_*mfr*_. Complete lines describe the possible progressions between states whereas dotted lines describe the parasite transmission between humans and mosquitoes. Red, gray, and black lines display the flows of resistant, sensitive, and recovered.

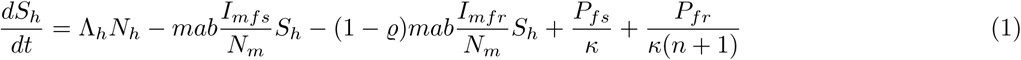

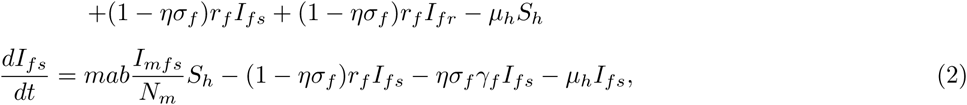

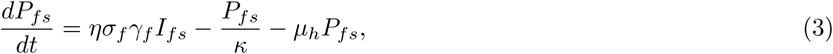

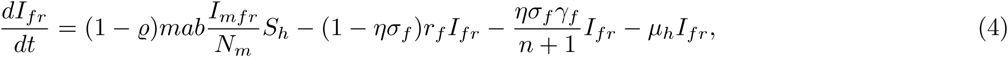

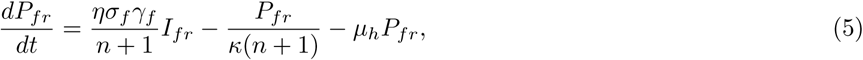

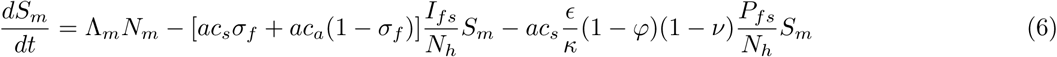

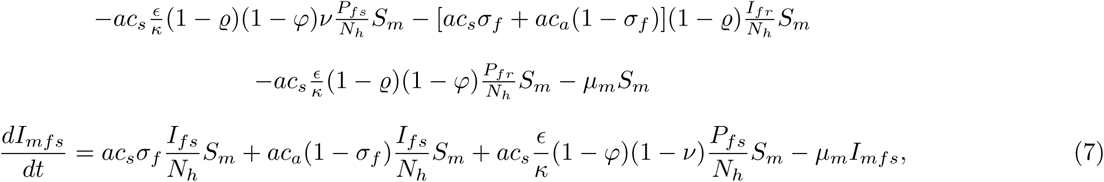

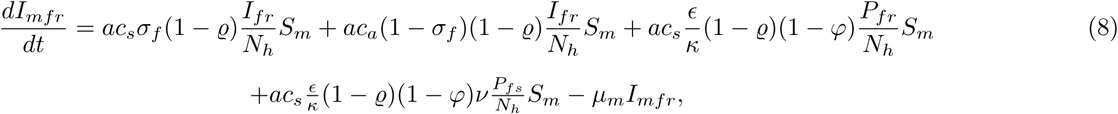

with

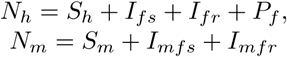

### *P. vivax* model

This model outlines *P. vivax* transmission in seven human and three mosquito states: susceptible humans *S*_*h*_, infected humans by sensitive strain *I*_*vs*_, humans with latent parasites of sensitive strain *L*_*vs*_, post-treatment humans after sensitive infection *P*_*vs*_, infected humans by resistant strain *I*_*vr*_, humans with latent parasites of resistant strain *L*_*vr*_, post-treatment humans after resistant infection *P*_*vr*_, susceptible mosquitoes *S*_*m*_, infected mosquitoes by sensitive strain *I*_*mvs*_, and infected mosquitoes by resistant strain *I*_*mvr*_. This model reproduces the same transmission interactions of *P. falciparum* model but involves two additional states: *L*_*vs*_ and *L*_*vr*_. These states depict humans with dormant hypnozoites of *P. vivax* that cause relapses after first infection. In fact, *I*_*vs*_, *I*_*vr*_, *P*_*vs*_ and *P*_*vr*_ can remain with latent parasites becoming *L*_*vs*_ or *L*_*vr*_ instead susceptible. Additionally, the model allows new infections in humans with latent parasites as Fig 2 illustrates. The equations are from the Eq 9 to Eq 18 using the parameters in table 1.

**Fig 2.**
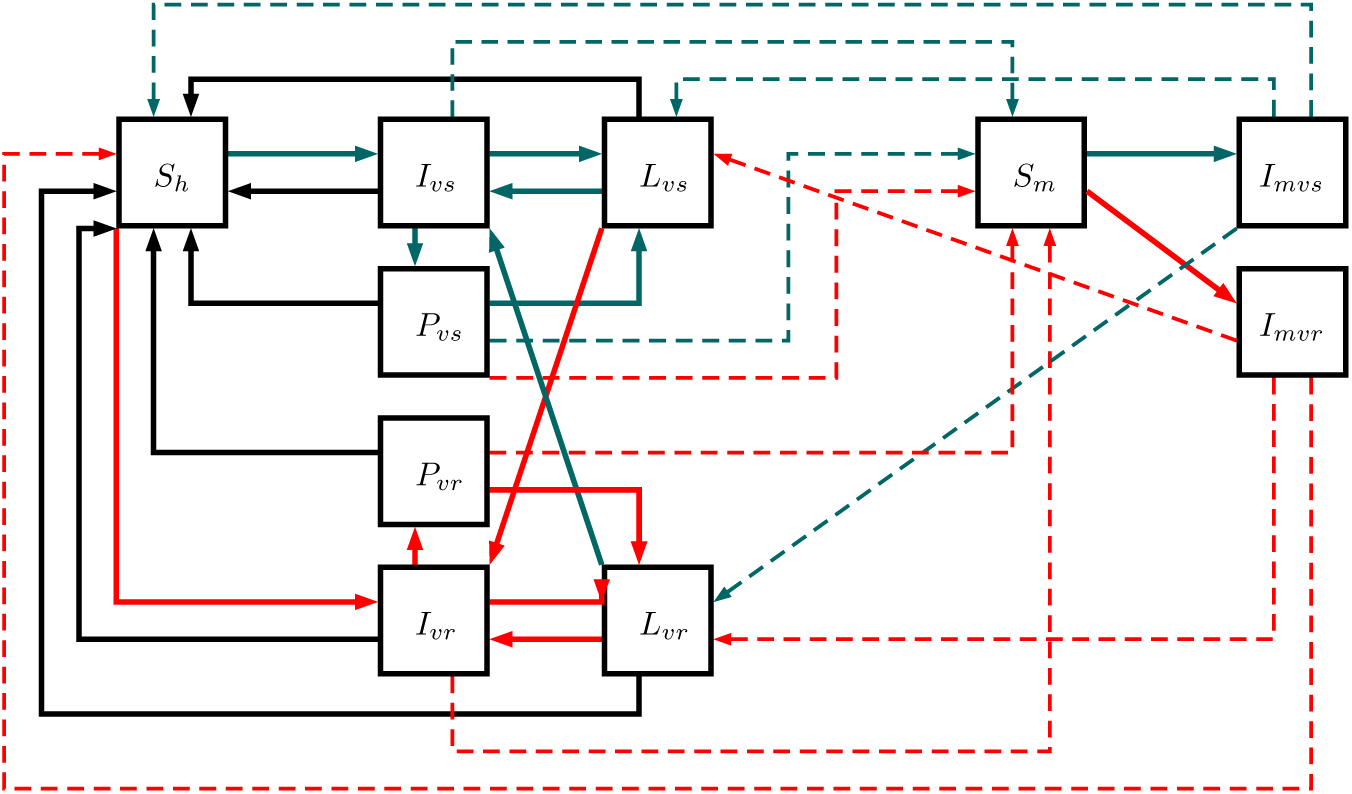
*P. vivax* model. This structure illustrates the transmission in seven human and three mosquito states: susceptible humans *S*_*h*_, infected humans by sensitive strain *I*_*vs*_, humans with latent parasites of sensitive strain *L*_*vs*_, post-treatment humans after sensitive infection *P*_*vs*_, infected humans by resistant strain *I*_*vr*_, humans with latent parasites of resistant strain *L*_*vr*_, post-treatment humans after resistant infection *P*_*vr*_, susceptible mosquitoes *S*_*m*_, infected mosquitoes by sensitive strain *I*_*mvs*_, and infected mosquitoes by resistant strain *I*_*mvr*_. Complete lines reproduce the possible progressions between states while dotted lines reproduce the parasite transmission between humans and mosquitoes. Red lines display the flow of resistant parasites, gray lines display the flow of sensitive parasites, and black lines display the flow without parasites.

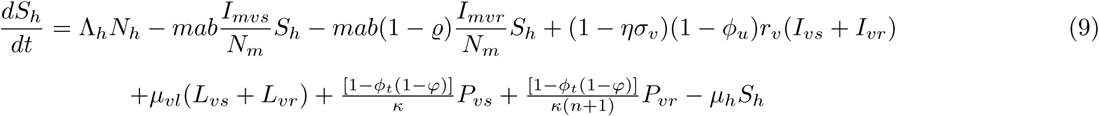

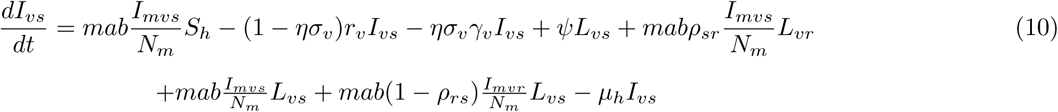

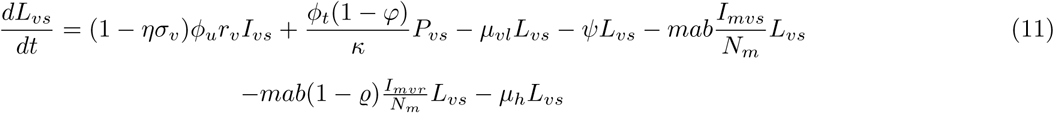

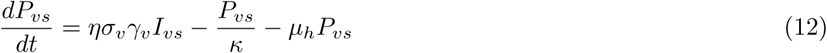

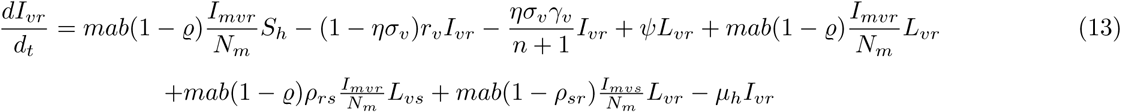

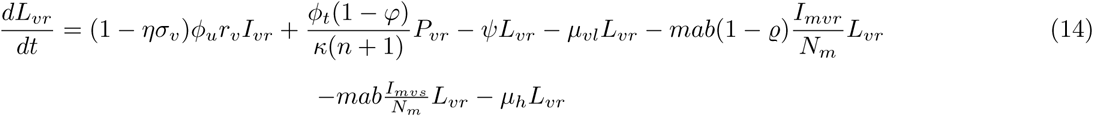

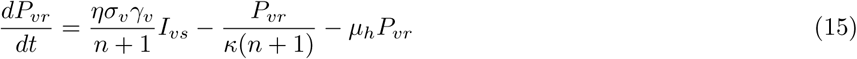

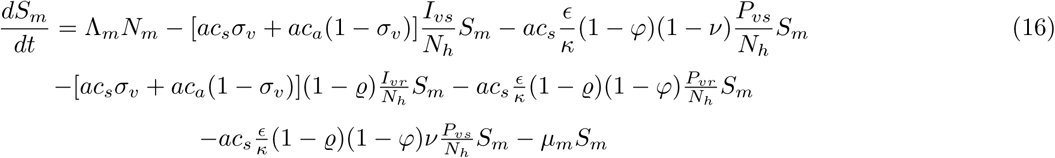

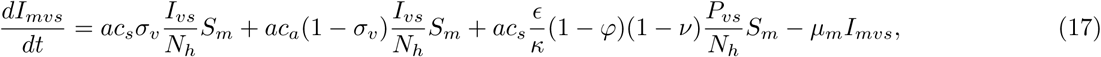

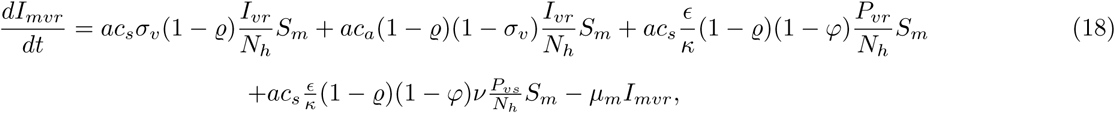

with

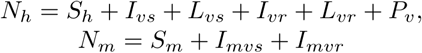

### Resistance cost

Resistance cost (*ρ*) reduces parasite fitness when a mutation occurs and generates resistance against a specific treatment [13]. We modeled this cost as a transmission reduction in resistant strains multiplying the transmission rate with the term (1 *- ρ*) that decreases the transmission in a *ρ* percent.

### Asymptomatic infections

We considered asymptomatic humans as proportion of infected humans that do not search for antimalarial treatment. They act as parasite reservoirs but their transmission potential are lower than symptomatic humans. In the model, the transmission probabilities from asymptomatic and symptomatic individuals to susceptible mosquitoes occur with different probabilities *c*_*a*_ and *c*_*s*_, respectively, considering *c*_*a*_ *< c*_*s*_ [36]. This species–dependent parameter is a consequence of the immunological profile in an endemic region due to exposition periods and it also varies according with *Plasmodium* specie [28]. Hence, we considered (1 − *σ*_*f*_) and (1 − *σ*_*v*_) as constant proportion of asymptomatic humans infected by *P. falciparum* and *P. vivax* assuming a long exposition period before treatment.

### Antimalarial treatment

Treatment coverage *η* varies from 0% to 100% of infected humans, adopting a single treatment-regimen. Additionally, the model also permits an evaluation of treatment plus primaquine in by a *ϕ* proportion of treated humans impacting gametocyte transmission and the *P. vivax* hypnozoites.

### Infectious period

Infected without available treatment and asymptomatic humans recover from infection at *r* rate with 1*/r* as infectious period because they do not employ treatment. On the other hand, treated humans advance to post-treatment state at *γ* rate with 1*/γ* as infectious period. Moreover, 1*/γ*_*v*_ *>* 1*/γ*_*f*_ because the early development of gametocytes in *P. vivax* triggers longer infectious period before treatment than *P. falciparum* [28, 46].

Likewise, we differentiated the infectious periods between sensitive and resistant strains according with the recurrences associated at resistant parasites [37]. The mean infectious time for a sensitive strain is 1*/γ* infectious period, whereas mean infectious period of a resistant strain is (*n* + 1)*/γ*, with *n* recurrences. This is because humans infected by resistant parasite extend their infectious periods when a recurrence occurs.

### Post-treatment period

Post-treatment period engages three dynamics: parasite clearance, drug half-life and emergence of resistant parasites. Parasite clearance of drugs such as chloroquine and artemisinin components differs and also affects specific parasite forms per species [38, 39]. Parasite clearance *E* characterizes the infectious period after treatment. Drug half-life *κ* corresponds to time interval when treatment remains in the blood conferring a protective period [40]. The emergence of resistant parasites occurs by the selection of parasite strains under residual drug concentration and we defined *ν* as the probability of transmitting a resistant parasite from a post-treatment infected by a sensitive strain [11].

### Basic reproduction number

We derived the basic reproduction number adopting the next generation matrix approach proposed in [41–43]. We assumed constant populations in humans (*N*_*h*_) and mosquitoes (*N*_*m*_) where *Λ*_*h*_ = 0, *µ*_*h*_ = 0 and *Λ*_*m*_ = *µ*_*m*_. In this way, we employed the disease free disease-stable steady state (*S*_*h*_ = *N*_*h*_; *S*_*m*_ = *N*_*m*_; the rest states equal to 0) to linearize the equations and then, we build the transmission and transition matrix in order to derive the basic reproduction numbers.

### Simulation

Our aim is to simulate the spread of drug resistance in *P. falciparum* and *P. vivax* species comparing between common-employed treatment regimens. We contrasted regimens between the adoption of four treatment lines: chloroquine (CQ), chloroquine plus primaquine (CQ+PQ), artemisinin combination therapy (ACT) and ACT plus PQ (ACT+PQ). The initial condition is only the presence of the sensitive strain and the table 2 summarized the parameters to each treatment regiment. We implemented the equations in RStudio using deSolve package [44].

**Table 2.**
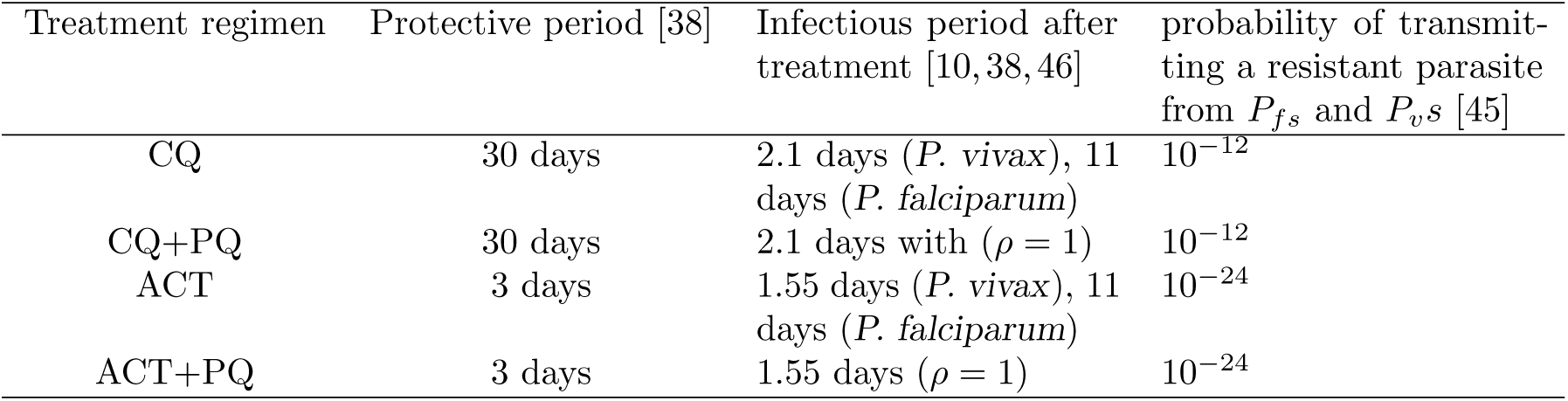
Treatment parameters.

| Treatment regimen | Protective period [38] | Infectious period after treatment [10, 38, 46] | probability of transmitting a resistant parasite from $P_{fs}$ and $P_{vs}$ [45] |
| --- | --- | --- | --- |
| CQ | 30 days | 2.1 days (*P. vivax*), 11 days (*P. falciparum*) | $10^{-12}$ |
| CQ+PQ | 30 days | 2.1 days with ($\rho = 1$) | $10^{-12}$ |
| ACT | 3 days | 1.55 days (*P. vivax*), 11 days (*P. falciparum*) | $10^{-24}$ |
| ACT+PQ | 3 days | 1.55 days ($\rho = 1$) | $10^{-24}$ |

### Sensitivity analysis

Finally, we performed a sensitivity analysis of parameter models on the emergence-time of the resistant strain using Latin Hypercube Sampling (LHS) to respond at the uncertainty of estimated values and also to assess the parameter influence [47]. We implemented the analysis in RStudio using deSolve, lhs and sensitivity packages [44, 48, 49].

## Results

### Basic reproduction number

We found the basic reproduction number for the sensitive and resistant strains in both *P. falciparum* and *P. vivax* models (see from Eq 19 to Eq 22). Resistance cost cuts down *R*_0_s of resistant strains in comparison with sensitive while the recurrences increase them for both species. On the other hand, terms associated with latent *vivax* parasites reduce *R*_0_ of this specie in same proportion between strains.

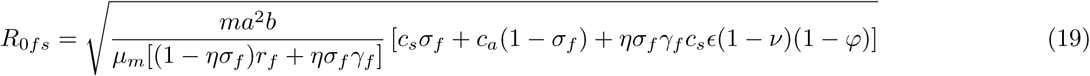

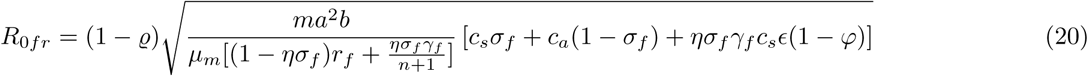

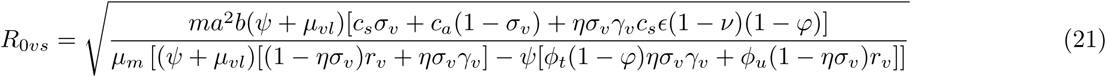

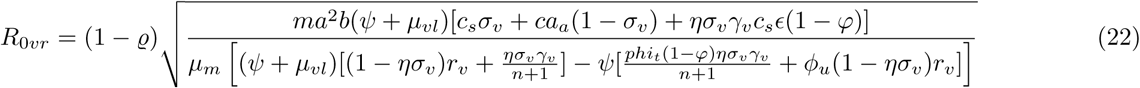

We evaluated the basic reproduction number *R*_0_ by varying cost resistance, treatment plus primaquine and infectious time before and after treatment (see Fig 3). In general, increases in drug coverage decrease *R*_0_s of *P. falciparum* at higher rate than *R*_0_s of *P. vivax. R*_0_ of sensitive *P. vivax* overcomes more *R*_0_s of resistant stains at different resistance cost than *R*_0_ of sensitive *P. falciparum*. Treatment plus primaquine influences at same the *R*_0_s of sensitives and resistant stains for both species. A longer infectious period before and after treatment induces highest *R*_0_s but only the longer infectious period before boosted the sensitive *R*_0_ to stay over the resistant *R*_0_; this effect is higher in *P. vivax*.

**Fig 3.**
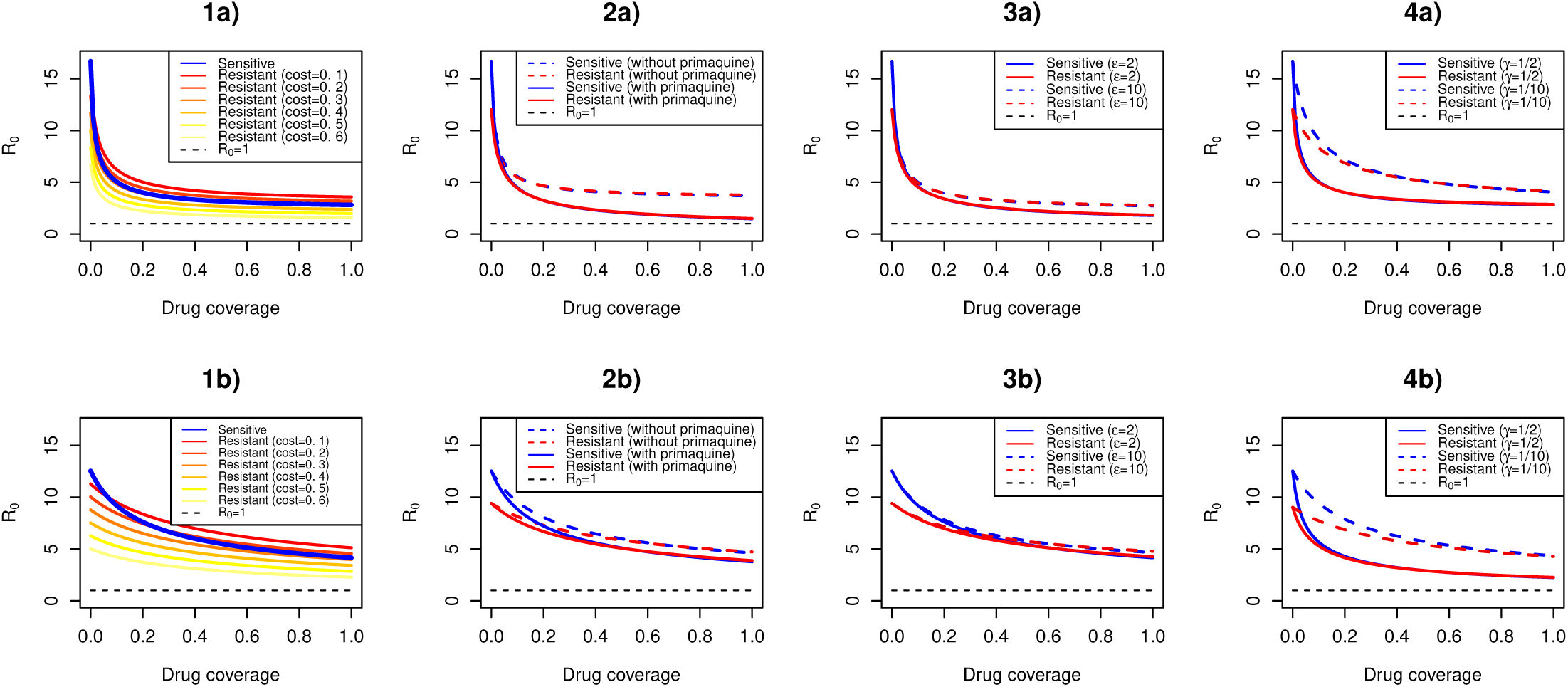
Drug coverage varying the basic reproduction numbers. The figure illustrates *R*_0_ lines for *P. falciparum* (figures a) and *P. vivax* (figures b) models dividing by sensitive and resistant strains. 1(a and 1(b display *R*_0_ lines of sensitive and resistant strains with different resistance cost; 2(a and 2(b display *R*_0_ lines using or non-using primaquine; 3(a and 3(b display *R*_0_ lines at two infectious periods after treatment in days (*ϵ*); 4(a and 4(b display *R*_0_ lines at two infectious periods before treatment in days (1*/γ*).

### Simulation

Although the regimens generated a more delayed emergence of resistant *P. vivax* strain, they accomplished a lesser reduction in the proportion of infected humans for this specie (see Fig 4). Treatment with chloroquine (CQ) contributed to a higher proportion of post-treatment humans, especially in the case of *P. falciparum*, and the emergence of resistant *P. vivax* took 2-fold as long time as resistant *P. falciparum*. Primaquine addition (CQ+PQ) decreased infected and post-treatment humans of *P. falciparum*, and humans with latent parasites of *P. vivax* but this regimen implied the emergence of resistant parasites in less time.

**Fig 4.**
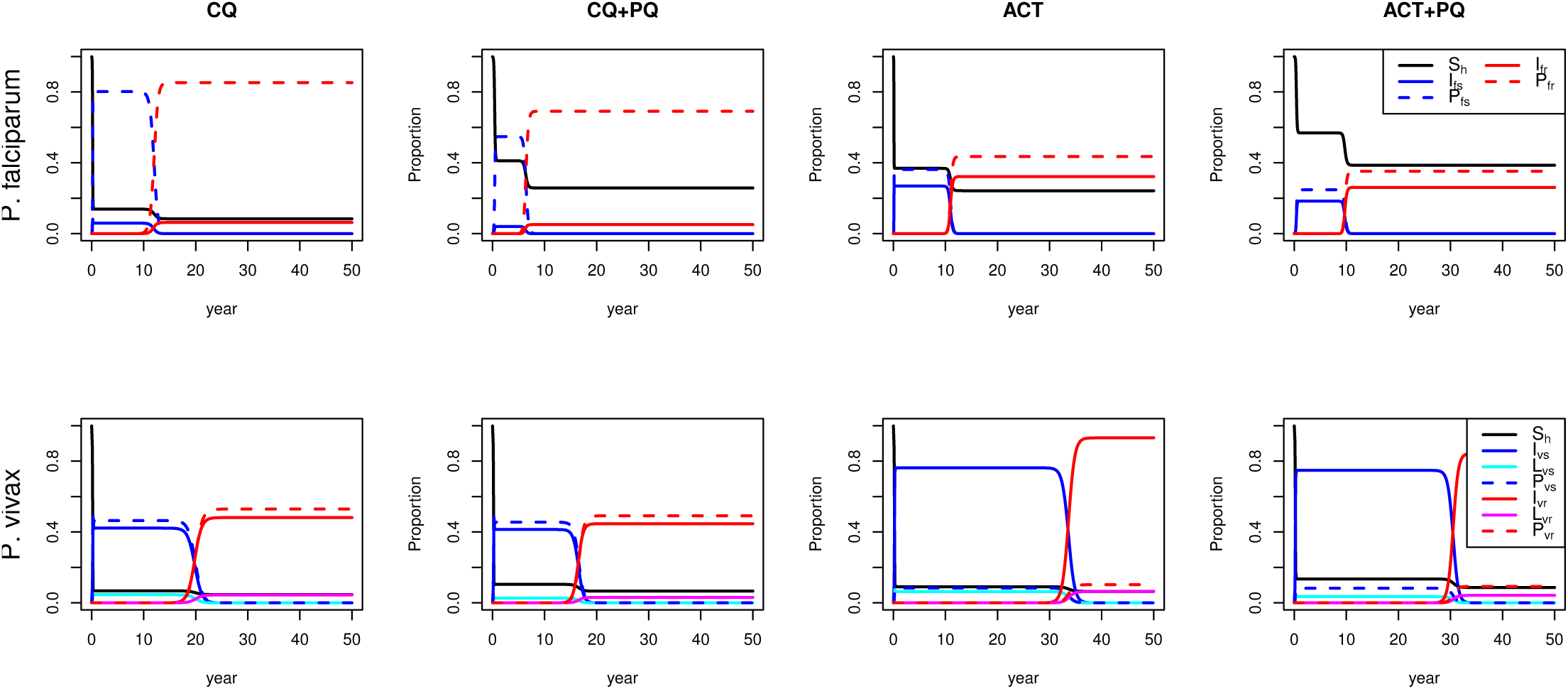
Simulation of treatment regimens. This figure illustrates the implementation of four treatment-regimens: chloroquine (CQ), chloroquine plus primaquine (CQ+PQ), artemisinin combination therapy (ACT) and artemisinin combination therapy plus primaquine (ACT+PQ). First row shows the simulated regimens in *P. falciparum* model and second row shows the simulated regimens in *P. vivax* model.

Regimens with artemisinin combination therapy delayed the emergence of resistant *P. vivax* three times as long as resistant *P. falciparum* but this regimen affected the proportion of infected humans less than chloroquine regimen. Primaquine addition (ACT+PQ) also decreased infected and post-treatment humans of *P. falciparum*, and humans with latent parasites of *P. vivax* though the emergence of resistant parasites remained at a similar time as the ACT without primaquine.

### Sensitivity analysis

In this model, the maximum influence is given by resistance cost with a proportional relation which implies this parameter is the one that more delays the emergence of resistant parasites either *P. vivax* or *P. falciparum* (see Fig 5). Five parameters also obtained proportional relation but with a low parameter influence: transmission probability from an symptomatic to susceptible mosquito, infectious period of post-treatment human, probability of developing sensitive infection by the contact between *I*_*mvs*_ and a *L*_*vs*_ (only in *P. vivax*) and probability of post-treatment human of remaining with latent parasites (only in *P. vivax*).

**Fig 5.**
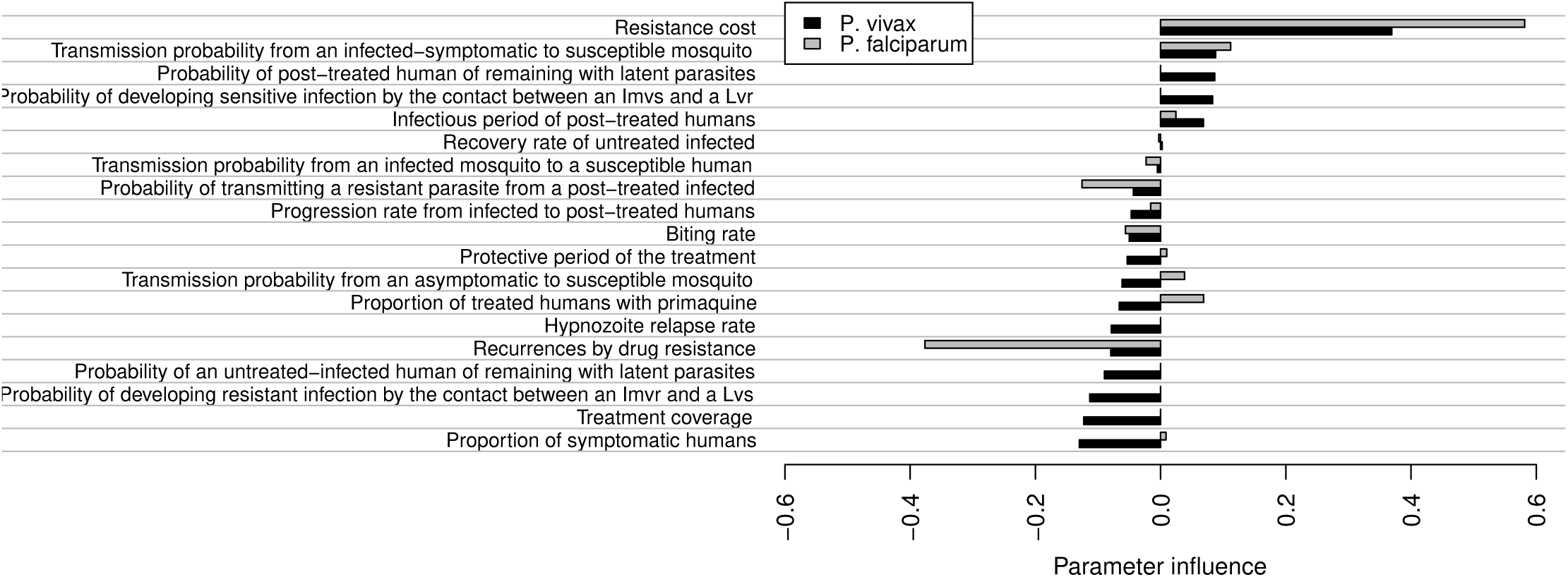
Parameter sensitivity on the emergence-time of the resistant strain. The figure illustrates parameter influence where -1 represents the maximum inverse relation (accelerate drug resistance), 1 represents the maximum proportional relation (delay drug resistance) and 0 represents no relation.

Recurrences by drug resistance obtained the most negative influence in *P. falciparum* that implies an earlier emergence of the resistant strain with increments in this parameter. Ten parameters also obtained a negative influence but with a lower value (see Fig. 5). On the other hand, three parameters had negative influence in *P. vivax* and positive in *P. falciparum* but with a lower value: transmission probability from an asymptomatic to susceptible mosquito, proportion of treated humans with primaquine and the protective period.

## Discussion

We found that the early transmission before treatment and the low effectiveness of drug coverage support the prevalence of sensitive parasites delaying the emergence of resistant *P. vivax*. In fact, the reproduction numbers of sensitive *P. vivax* surpassed the reproduction numbers of resistant ones when the infectious period before treatment was greater and this usually occurs in *P. vivax* transmission by the early development of gametocytes [28]. On the other hand, all treatment regimens exhibited a lower response against the proportion of *P. vivax* infected-humans. This commonly happens with control efforts that work slowly against *P. vivax* [50].

Artemisinin-based combinated therapy (ACT) and Chloroquine (CQ) caused the emergence of *P. falciparum*-resistant in a similar time scale whereas ACT delayed the emergence of *P. vivax*-resistant. In theory, development of drug resistance to a set of drugs is less likely than a single drug and this reinforces the improvement of combined therapies [45].

Nevertheless, our results show that the rapid parasite-clearance and the shorter protective period with ACT against *P. falciparum* avoided the transmission of sensitive parasites after-treatment allowing a similar emergence period of resistant parasite as CQ in spite of the lower probability of transmitting resistant parasites with ACT. Actually, ACT-resistance in *P. falciparum* have emerged after 10 years of use in 2008 while CQ-resistance emerged in the 50’s, ten years after CQ adoption, indicating a connection with the current results [51, 52].

*P. vivax* CQ-resistant emerged earlier than ACT resistant but CQ achieved a higher reduction in the prevalence of infected humans in overall simulations. Previous studies reported CQ-resistance whereas we have no reports of ACT-resistance in *P. vivax* and our simulated ACT-regimen exhibited a longer emergence of resistant strain than the period since ACT adoption bringing an explanation of the no reported ACT-resistance [53]. On the other hand, treatment plus primaquine (PQ) declined the proportion of humans with latent parasites but it accelerated the emergence of resistant strain because PQ avoids the transmission of sensitive parasites after-treatment. This result challenges the suggestion of PQ-dose to avoid the transmission of resistant parasites and *P. vivax* relapses can explain the underestimate resistance induced by PQ effect [53, 54].

The main limitation to validate the results and model parameterization is the necessary data in distinct ways: disease prevalence, resistance frequency, treatment effort and transmission conditions. Data bases such as the Malaria Atlas Project (MAP) and the World Wide Antimalarial Resistance Network (WWARN) provide reports of disease incidence and studies for monitoring the drug resistance but several factors causes disturbances to track drug resistance: different sample sizes in clinical trials, intermittent periods of monitoring, absent of drug-coverage measures and different levels of resistance [55, 56]. Therefore, we accomplished a sensitive analysis that revealed the resistance cost and the recurrences by drug resistance as the most influential parameters and the rest of parameters had lower influence on the emergence of resistant parasites.

*P. vivax* and *P. falciparum* develop drug resistance but the pace depends on the maintenance of sensitive parasites. In fact, *P. vivax* supported sensitive parasites through the transmission before treatment, reservoirs of latent parasites and asymptomatic humans causing a less effectiveness in treatment-regimen effort. On the other hand, *P. falciparum* develops an earlier resistance because treatment regimens accomplished a greater reduction in the reproduction numbers and proportion of infected humans in sensitive parasites. These results imply that early attention of symptomatic humans and reservoirs of *P. vivax* helps reduce the population of sensitive parasites but it would accelerate drug resistance.

## Acknowledgments

MA and DV are grateful for the Print internationalization program supported by Coordenação de Aperfeiçoamento de Pessoal de Nível Superior (CAPES, http://www.capes.br) and Fundação Oswaldo Cruz (Fiocruz, http://www.fiocruz.br) and MA is grateful for the scholarship support from Instituto Oswaldo Cruz (IOC, http://www.ioc.fiocruz.br) in the graduate program. DV has support from Conselho Nacional de Desenvolvimento Científico e Tecnológico (CNPq, http://www.cnpq.br, projeto Universal, Ref. 424141/2018-3).

